# KAKU4-mediated deformation of the vegetative nucleus controls its precedent migration over sperm cells in pollen tubes

**DOI:** 10.1101/774489

**Authors:** Chieko Goto, Kentaro Tamura, Satsuki Nishimaki, Naoki Yanagisawa, Kumi Matsuura-Tokita, Tetsuya Higashiyama, Daisuke Maruyama, Ikuko Hara-Nishimura

## Abstract

A putative nuclear lamina protein, KAKU4, modulates nuclear morphology in *Arabidopsis thaliana* seedlings but its physiological significance is unknown. *KAKU4* was strongly expressed in mature pollen grains, each of which has a vegetative cell and two sperm cells. KAKU4 protein was highly abundant on the envelopes of vegetative nuclei (VNs) and less abundant on the envelopes of sperm cell nuclei (SCNs) in pollen grains and elongating pollen tubes. VN is irregularly shaped in wild-type pollen. However, *KAKU4* deficiency caused it to become more spherical. These results suggest that the dense accumulation of KAKU4 is responsible for the irregular shape of the VNs. After a pollen grain germinates, the VN and SCNs migrate to the tip of the pollen tube. In the wild type, the VN preceded the SCNs in 91–93% of the pollen tubes, whereas in *kaku4* mutants, the VN trailed the SCNs in 39–58% of the pollen tubes. *kaku4* pollen was less competitive than wild-type pollen after pollination, although it had an ability to fertilize. Taken together, our results suggest that controlling the nuclear shape in vegetative cells of pollen grains by *KAKU4* ensures the orderly migration of the VN and sperm cells in pollen tubes.

**Highlight:** The nuclear envelope protein KAKU4 is involved in controlling the migration order of vegetative nuclei and sperm cells in pollen tubes, affecting the competitive ability of pollen for fertilization.

## Introduction

In *Arabidopsis thaliana*, pollen grains contain a vegetative cell and two sperm cells (Borg and Twell, 2011; McCue *et al.*, 2011). One of the sperm cells fertilizes the egg cell to produce the embryo and the other fertilizes the central cell to produce the endosperm (Hamamura *et al.*, 2011; Kawashima and Berger, 2011; Maruyama *et al.*, 2015). The vegetative nucleus (VN) has irregular shapes (Vogler *et al.*, 2015) and is physically connected to one of the two sperm cells. Together, the VN and the two sperm cells form a functional unit termed the male germ unit (McCue *et al.*, 2011). After pollination, the pollen grain germinates. Then, the VN and its trailing sperm cells move into and migrate along the elongating pollen tube, although they occasionally change their migration order (Zhou and Meier, 2014).

Nuclear movements are driven by the plant-specific motor protein myosin XI-i (Tamura *et al.*, 2013). In *A. thaliana*, the migration order of the VN and sperm cells is not affected by myosin XI-I deficiency (Zhou and Meier, 2014), but it is affected by the outer nuclear membrane proteins WIPs (WIP1, WIP2, and WIP3) and WITs (WIT1 and WIT2) and the possible inner nuclear membrane proteins SUNs (SUN1 and SUN2) (Zhou *et al.*, 2015; Zhou and Meier, 2014).

Plant nuclei have a fibrillar meshwork lamina structure beneath the inner nuclear membrane (Fiserova *et al.*, 2009). KAKU4 (after the Japanese word for nucleus, *kaku*) is a candidate component of the plant nuclear lamina (Goto *et al.*, 2014; Meier *et al.*, 2017; Poulet *et al.*, 2017). In epidermal cells of the mature vegetative tissues of *A. thaliana*, a KAKU4-deificient mutant (*kaku4*) has smaller spherical-shaped nuclei, compared with spindle-shaped nuclei of the wild type. A spherical phenotype is also found in mutants defective in the outer nuclear membrane proteins WIPs (Zhou *et al.*, 2012), in the possible inner nuclear membrane proteins SUNs (Zhou *et al.*, 2012), and in the nuclear envelope-associated proteins CRWNs (CRWN1 and CRWN4) (Dittmer *et al.*, 2007; Goto *et al.*, 2014; Sakamoto and Takagi, 2013; Wang *et al.*, 2013).

In this study, we show that *KAKU4* is highly expressed in the VNs of pollen grains and tubes, resulting in deep invagination of the VN envelope. We propose that KAKU4-mediated deformation of the VN controls the proper migration order of VN and sperm cells during pollen tube growth in *A. thaliana*.

## Materials and methods

### Gene expression data from public databases

We assessed the gene expression level data in various tissues deposited in the database Atted-II (http://atted.jp/), and then downloaded the folder “Expression_data_Development” and used the file “AtGE_dev_gcRMA” which contains the raw data of microarray experiments (Schmid *et al.*, 2005). We chose the values of the genes of interest from wild-type plants for graphical presentation.

### Plant materials

*Arabidopsis thaliana* (Columbia-0) was used as the wild-type line. T-DNA insertion mutants (SALK_076754 [*kaku4-2*], SALK_010298 [*kaku4-3*], SAIL_711_E09 [*kaku4-4*], and SALK_041774 [*crwn1-2*],) and EMS-mutagenized mutants (*kaku2* and *kaku4-1*) were isolated previously (Goto *et al.*, 2014). The transgenic plants *ProKAKU4:KAKU4-tRFP kaku4-2* and *ProKAKU4:KAKU4-EYFP kaku4-3*, in which the genomic fragment of the splice variant *KAKU4.2* was expressed, were also used (Goto *et al.*, 2014). Seeds were germinated and grown on MS medium containing 0.5% gellan gum at 22°C under continuous light (35 µmol m^−2^ s^−1^). The plants were transferred to vermiculite in 2-3 weeks and grown at 22°C under a light cycle of 16 h light/8 h dark.

### Plasmid construction and transformation using GUS under the KAKU4 promoter control

To construct *ProKAKU4:GUS*, a genomic fragment encompassing the 2-kb sequence upstream of the coding sequence was cloned into pENTR/D-TOPO (Invitrogen) and then fused upstream of *EGFP-GUS* in a plant transformation vector (pKGWFS7). The primers used for the amplification of the *KAKU4* promoter were the forward primer (5′-CACCGAGGAATACAGGCGAGAACA-3′) and the reverse primer (5′-GGTGAAAGTGAGAGGAGGAG-3′). The sequence motif 5′-CACC-3′ required for cloning into pENTR/D-TOPO was added at the 5′-end of the forward primer. To obtain stable expression, *Arabidopsis* wild-type plants were transformed with *Agrobacterium tumefaciens* (GV3101) using the floral dip method (Clough and Bent, 1998).

### GUS staining

Floral organs were immersed in 90% acetone, and stored for one or two nights at −20°C. Then, 90% acetone was replaced by 1× phosphate buffer (100 mM NaPO_4_; 10× stock solution [1M NaPO_4_] was prepared by mixing 1M Na_2_HPO_4_ and 1M NaH_2_PO_4_ at a ratio of 61:39; 10× stock solution was diluted 10 times and used as 1× phosphate buffer), which was immediately replaced by the GUS-staining solution (100 mM NaPO_4_ [pH 7.4], 10 mM EDTA, 0.1% Triton X-100, 1 mM potassium ferricyanide, 1 mM potassium ferrocyanide, 0.5 mg/mL 5-bromo-4-chloro-3-indolyl-β-D-glucuronide cyclohexylammonium). The GUS-staining solution was vacuum infiltrated into the floral organs for 5 min at 20-25°C. The floral organs infiltrated by the GUS-staining solution were incubated for 16 h in the dark at 37°C for GUS-staining reaction. The GUS-staining solution was replaced by 1× phosphate buffer, followed by rinsing with 70% ethanol two times. After rinsing, an ethanol and acetic acid mixture (ethanol: acetic acid = 6:1) was added for specimen preparation. The specimens were mounted with chloral hydrate solution (chloral hydrate: glycerol: water = 8 [g]: 1 [ml]: 2 [ml]) and examined using a differencial interference contrast microscope (DIC) (Axioskop 2 plus system [Carl Zeiss] and a high-sensitivity cooled CCD color camera VB-7010 [Keyence]).

### Microscopy

Confocal fluorescence images except for those in Fig. 6 were obtained using a confocal laser scanning microscope (LSM780; Carl Zeiss). The 405 nm line of a blue diode laser, the 488 nm line of a 40 mW Ar/Kr laser, and the 544 nm line of a 1 mW He/Ne laser were used to excite Hoechst 33342, GFP/YFP, and RFP, respectively. Images were acquired with a ×63 1.2 NA water immersion objective (C-Apochromat, 441777-9970-000; Carl Zeiss), a ×40 0.95 NA dry objective (Plan-Apochromat, 440654-9902-000; Carl Zeiss), or a ×20 0.80 NA dry objective (Plan-Apochromat, 440640-9903-000; Carl Zeiss). Data were exported as TIFF files and processed using Adobe Photoshop Elements 9.0 (Adobe Systems) or ImageJ 1.45s (National Institutes of Health; NIH). Fluorescence images presented in Fig. 5A were obtained using an Axioskop2 plus microscope (Carl Zeiss) and a high-sensitivity cooled CCD color camera VB-7010 (Keyence).

### Hoechst 33342 staining

Pollen grains and pollen tubes were stained with a solution containing 1 µg/ml Hoechst 33342, 3.7% (w/v) paraformaldehyde, 10% (v/v) dimethyl sulfoxide, 3% (v/v) Nonidet P-40, 50 mM PIPES-KOH (pH 7.0), 1 mM MgSO_4_, and 5 mM EGTA.

### *In vitro* pollen germination

*In vitro* pollen germinations were performed as described previously (Boavida and McCormick, 2007). Briefly, pollen grains from full-open flowers were transferred to the medium on a glass slide by gently scraping the surface of the medium by a flower or anthers. Three or four glass slides with the medium containing transferred pollen were placed on Kim Wipes (produced by Nippon Paper Crecia based on partnership with Kimberly Clark) in a square-shaped plastic petri dish (EIKEN CHEMICAL). Kim Wipes wetted with water were put along the vertical wall of the dish to enclose the glass slides before covering the dish with the lid. The petri dish was placed in an incubator with temperature control set to 22°C and continuous light for 3–3.5 h, followed by microscopic observation.

### Generation of double nuclear marker line for *in vitro* pollen tube growth assay

pDM441, a binary vector harboring *ProLAT52:NLS-Clover* was generated as follows. DNA fragment encoding mClover with A206K was amplified from a template plasmid pPZP221 CloN (Takeuchi and Higashiyama, 2016) by a PCR using a primer set pENTR_NLS_GFP (5’-CAC CAT GGC TCC AAA GAA GAA GAG AAA GGT CAT GGT GAG CAA GGG CGA GGA G −3’) and GFP_R (5’-CTT GTA CAG CTC GTC CAT GCC G −3’). The PCR product was cloned into the pENTR/D-TOPO (Thermofisher) to generate pOR003. LR recombination was performed between the pOR003 with a modified pGWB501 (Nakagawa *et al.*, 2007) harboring *LAT52* promoter at the *Hin*dIII site (Twell *et al.*, 1990) to produce the pDM441. Agrobacterium GV3101 containing the pDM441 or *ProRPS5A:HISTONE 2B-tdTomato* (Maruyama *et al.*, 2013) were independently cultured and subsequently used in simultaneous transformation by the floral dip method (Clough and Bent, 1998) to generate double nuclear marker line (*ProRPS5A:H2B-tdTomato ProLAT52:NLS-Clover*).

### Analysis of nuclear size and shape in pollen tubes in the microfluidic device

Pollen grains from the *ProRPS5A:H2B-tdTomato ProLAT52:NLS-Clover* double nuclear marker line were suspended in an *in vitro* pollen tube growth medium (*Muro et al., 2018*) without agarose and placed in a polydimethylsiloxane (PDMS) microfluidic device on a glass slip (Matsunami). The PDMS microfluidic device with 10 µm-width of micro channels was prepared as described previously (Yanagisawa *et al.*, 2017). The device allows a pollen tube to elongate on a focal plane, resulting in easy imaging analysis for hours. After 1h incubation at 22 ºC, fluorescence images of the nuclei labeled with H2B-tdTomato and NLS-Clover in pollen tubes were obtained using a confocal laser scanning microscope SP8 (Leica Microsystems). The images were exported as TIFF files and processed using the Analyze Particles function of ImageJ. A two-tailed homoscedastic Student’s t test was performed using Microsoft Excel. The circularity index was calculated using the equation 4πA/P^2^ (where A = area of nucleus and P = perimeter of nucleus) and indicates how closely each nucleus corresponds to a spherical shape (a perfect sphere has a circularity index of 1). Any deviation from a circular shape (e.g., elongated, lobulated, or spindle shaped) causes the index to decrease.

### Clearing siliques

Fixation and clearing of siliques were performed as described previously (Chen *et al.*, 2015). Silique length was measured using ImageJ software.

### Reciprocal crosses

F1 seeds from crosses between wild-type plants and *kaku4-3* hetero mutants (*kaku4-3*/+) were sown on MS medium containing kanamycin sulfate at 70 µg/ml. After 1-2 weeks, the live plants (Kan^R^) and dead plants (Kan^S^) were counted and subjected to chi-squared tests.

### Accession numbers

Sequence data from this article can be found in the Arabidopsis Genome Initiative or EMBL/GenBank databases under the following accession numbers: *CRWN1* (At1g67230), *CRWN2* (At1g13220), *CRWN3* (At1g68790), *CRWN4* (At5g65770), *KAKU4* (At4g31430), *Myosin XI-i* (At4g33200), *Nup136* (At3g10650), *SUN1* (At5g04990), *SUN2* (At3g10730), *WIP1* (At4g26455), *WIP2* (At5g56210), *WIP3* (At3g13360), *WIT1* (At5g11390), and *WIT2* (At1g68910).

## Results

### Remarkably high expression of *KAKU4* in mature pollen grains

By analyzing the gene-expression database of *A. thaliana* (ATTED-II, http://atted.jp/), we found that *KAKU4*, which encodes a putative nuclear lamina protein, is highly expressed in mature pollen (Fig. 1A) and that the high expression is specific to pollen (Supplemental Fig. S1). Similarly, *WIT1*, which encodes an outer nuclear membrane protein, is highly expressed in mature pollen (Fig. 1A) but it is also highly expressed in seeds (Supplementary Fig. S1). Expressions of *WIT2* and *WIP3*, which encode other nuclear membrane proteins, do not differ greatly among the tissues (Fig. 1A and Supplementary Fig. S1).

**Fig. 1.**
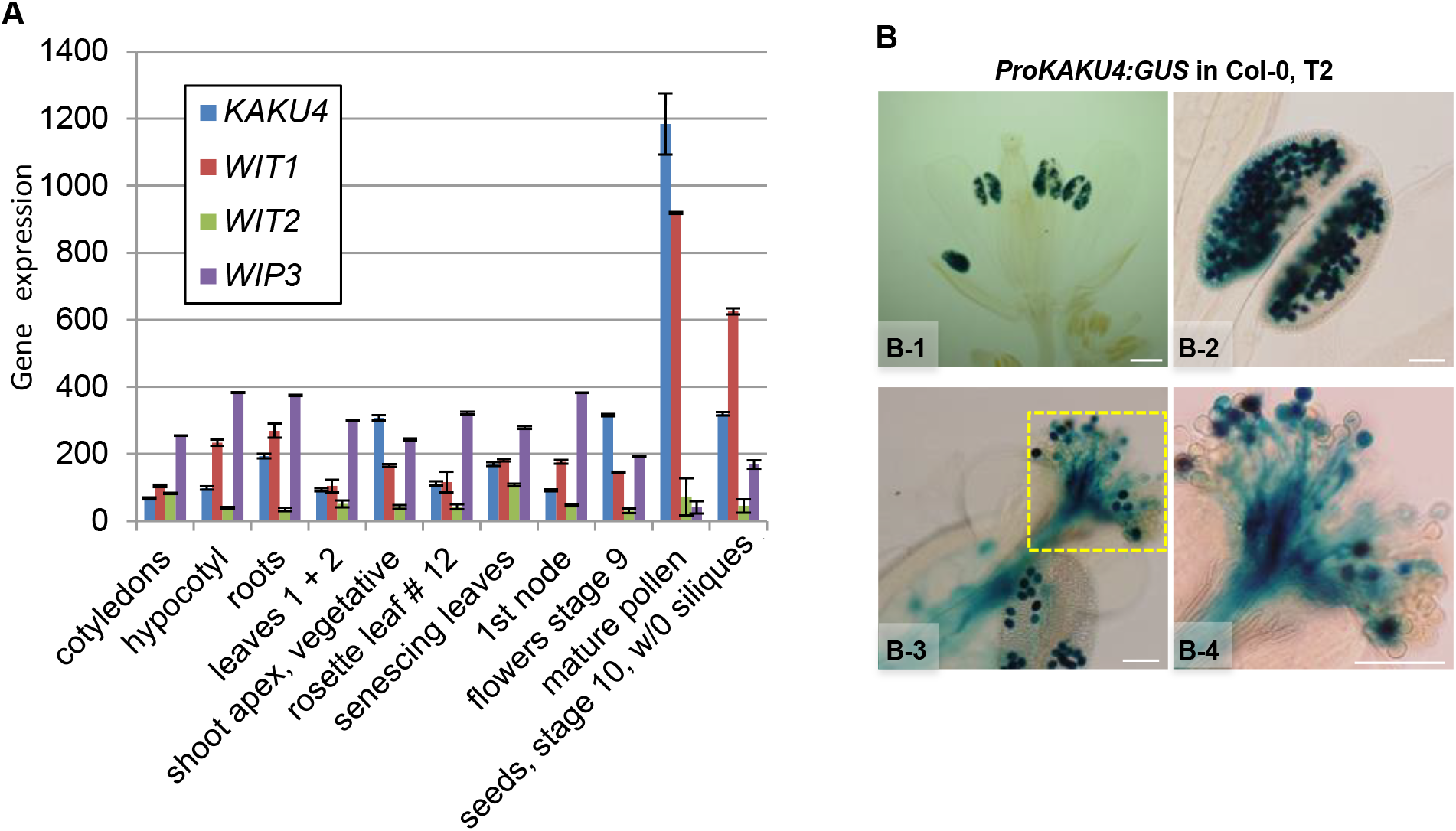
Expression of *KAKU4* in pollen. (A) Transcript levels of *KAKU4*, *WIT1*, *WIT2* and *WIP3* in various tissues. Data were obtained from the website of the AtGenExpress project (http://jsp.weigelworld.org/AtGenExpress/resources/) (Schmid et al. 2005). Intensities (absolute values) from some tissues (see the website for details) were extracted and the means ± SE are shown. The data for *WIP1* and *WIP2* were not available. See also Supplementary Fig.1 for the values from more tissues. (B) β-Glucuronidase (GUS) reporter assay of heterozygous transgenic plants that expressed GUS gene under the control of *KAKU4* promoter. (B-1) GUS-stained flowers. (B-2) Enlarged image of the anther. (B-3) Pollen attached to a stigma. (B-4) Enlarged image of the stigma in B-3. Some of the pollen grains are not stained because the tested line carries the heterozygous transgene (B-4). Scale bars = 500 µm in B-1 and 100 µm in B-2 to B-4.

To visualize the activity of the *KAKU4* promoter in mature pollen, we generated transgenic plants expressing the β-glucuronidase (GUS) reporter gene under the control of the *KAKU4* promoter. Strong GUS signals were detected in pollen grains of the anthers (Fig. 1B, upper panels) and pollen tubes elongating into the stigma (Fig. 1B, lower panels). These results indicate that *KAKU4* is predominantly expressed in mature pollen grains.

### KAKU4 is highly abundant on the VN envelope and less abundant on the SCN envelope in pollen grains

We examined the subcellular localization of the KAKU4 protein in mature pollen grains with the fluorescent KAKU4 markers. To avoid overexpression of the transgenes, KAKU4-tRFP and KAKU4-EYFP were expressed in the *KAKU4*-deficient mutant alleles *kaku4-2* and *kaku4-3*, respectively, under the control of the native promoter. Fluorescence signals of KAKU4-tRFP were clearly detected at the VN of each pollen grain (Fig. 2A). At a higher magnification, the fluorescence images showed that KAKU4-tRFP was localized to the nuclear envelope of the VN that were stained with Hoechst 33342 (Fig. 2B, upper panels). With an increased imaging sensitivity, KAKU4-tRFP was also detected on the envelope of two SCNs that were stained with Hoechst 33342 (Fig. 2B, lower panels, arrows). Similar results were obtained with KAKU4-EYFP (see Fig. 3A). The fluorescence levels of the SCNs were much lower than those of the VNs (Fig. 2B and Fig. 3A). Hence, KAKU4 is more densely accumulated in the envelopes of the VNs than in the envelopes of the SCNs.

**Fig. 2.**
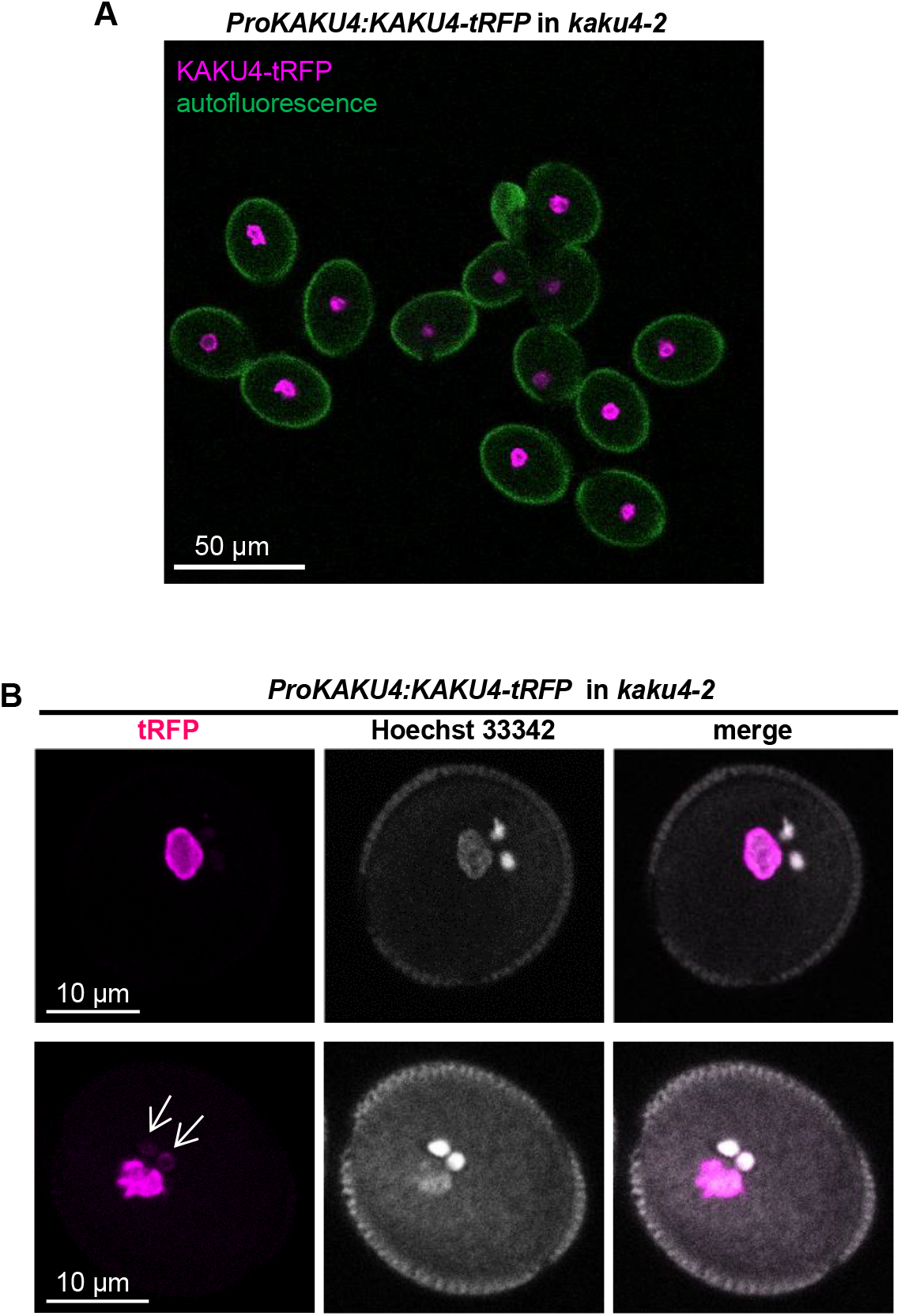
Subcellular localization of KAKU4 to the nuclear envelope in pollen grains. (A) Pollen grains from *kaku4-2* expressing *KAKU4-tRFP* under the control of *KAKU4* promoter. Magenta, tRFP; Green, autofluorescence. (B) Two sets of enlarged images of pollen grains from *kaku4-2* expressing *KAKU4-tRFP* (upper and lower). The pollen grains were stained with Hoechst 33342. Arrows indicate SCNs.

**Fig. 3.**
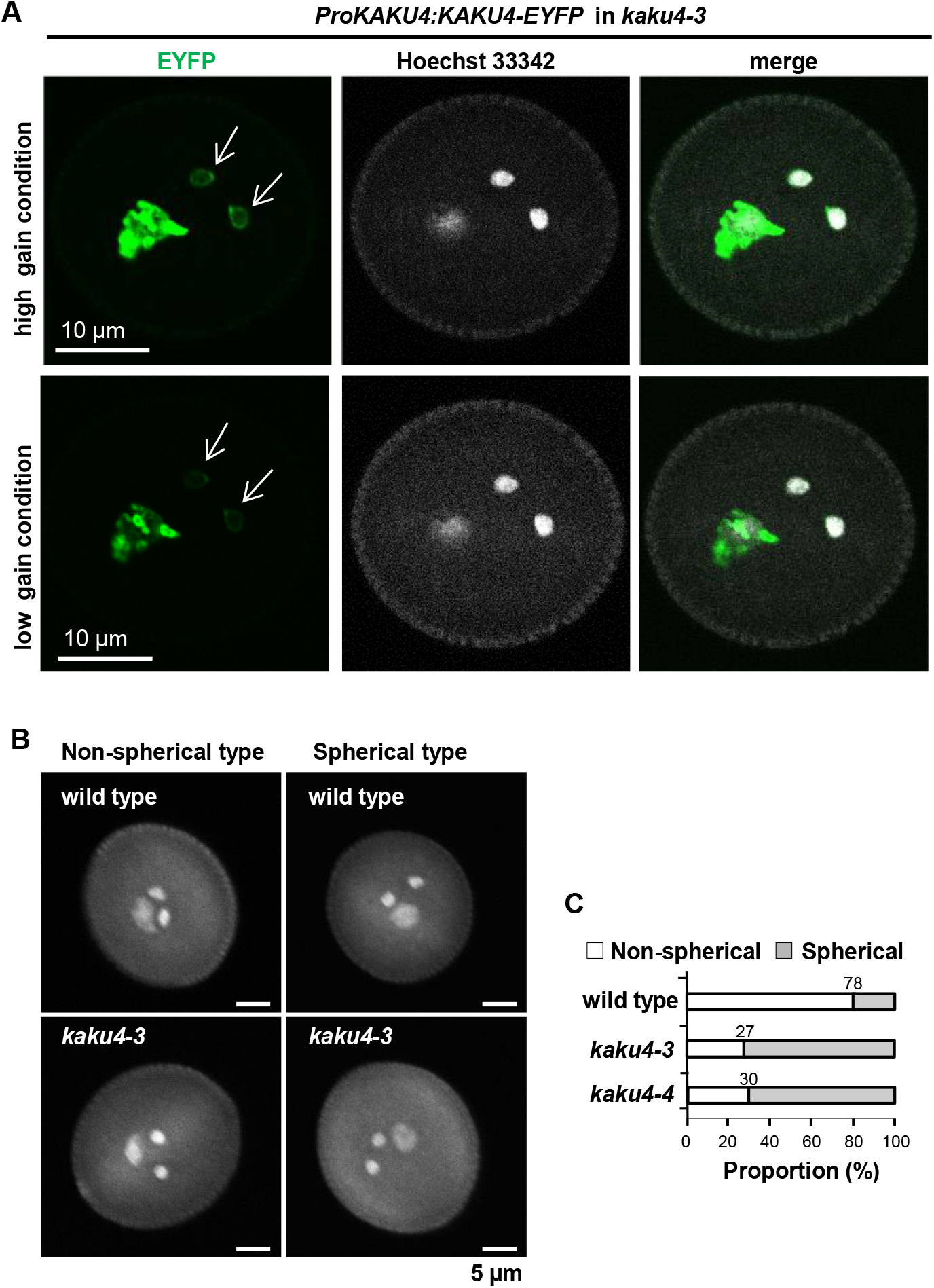
Nuclear shape of VN in pollen grains of *kaku4* (A) A pollen grain from *kaku4-3* expressing *KAKU4-EYFP* under the control of *KAKU4* promoter. Two sets of images with high and low digital gain (upper and lower panels, respectively) are shown. Arrows indicate SCNs. Note that the VN has invaginated nuclear envelope inside, while the SCNs have not. (B) Fluorescence images of nuclei stained with Hoechst 33342 in pollen grains. Non-spherical type, a pollen grain whose VN is irregularly shaped. Spherical type, a pollen grain of which VN is spherically shaped. (C) Proportions of VN with non-spherical shape and VN with spherical shape each for wild-type pollen and *kaku4* pollen. In total, 51 (wild type), 100 (*kaku4-3*), and 61 (*kaku4-4*) pollen grains were tested.

### The dense accumulation of KAKU4 leads to deformation of the VN

The nuclear shapes were compared between the SCN and the VN in the transgenic plants that expressed KAKU4-EYFP in *kaku4-3* under the control of the native promoter. The SCNs were nearly spherical in shape and their envelopes were relatively smooth (Fig. 3A). On the other hand, the VN were deeply invaginated (Fig. 3A). The deformation features of the VN are very similar to those of the nuclei in seedlings overexpressing KAKU4-GFP, in which artificial overexpression of KAKU4 causes nuclear envelope deformation in a dose-dependent manner (Goto *et al.*, 2014). This implied that the VN deformation is due to hyperaccumulation of KAKU4 on their envelopes.

Next, the effect of *KAKU4* deficiency on the deformation degrees of the VNs was examined with pollen grains from the wild type and the *kaku4* mutant alleles. The shapes of the VNs stained with Hoechst 33342 were classified into two types: spherical and non-spherical (Fig. 3B). Non-spherical shapes included elongated shapes and distorted shapes. The proportions of non-spherical shaped VNs were significantly lower in both *kaku4-3* and *kaku4-4* than in the wild type (Fig. 3C). Taken together, this result indicates that KAKU4 has a role in modulating nuclear shape in pollen.

### Involvement of KAKU4 in controlling the migration order of VN and SCNs in elongating pollen tubes

After pollination, a pollen grain germinates and elongates to form a pollen tube, in which the VN and the SCNs migrate towards the ovules (Kawashima and Berger, 2011). Using transgenic plants that expressed KAKU4-tRFP in *kaku4-2* under the native promoter and an *in-vitro* pollen germination assay (Boavida and McCormick, 2007), we were able to observe the KAKU4-labeled VN and the two SCNs migrating towards the tip of elongating pollen tubes (Fig. 4A, left). Like the nuclei in pollen grains, the VN envelopes highly fluoresced, while the SCN envelopes moderately fluoresced (Fig. 4A, right). Similar observations were made with another transgenic plant that expressed KAKU4-EYFP in *kaku4-3* under the native promoter (Fig. 4B). In most elongating pollen tubes, the VN preceded the two SCNs (Fig. 4).

**Fig. 4.**
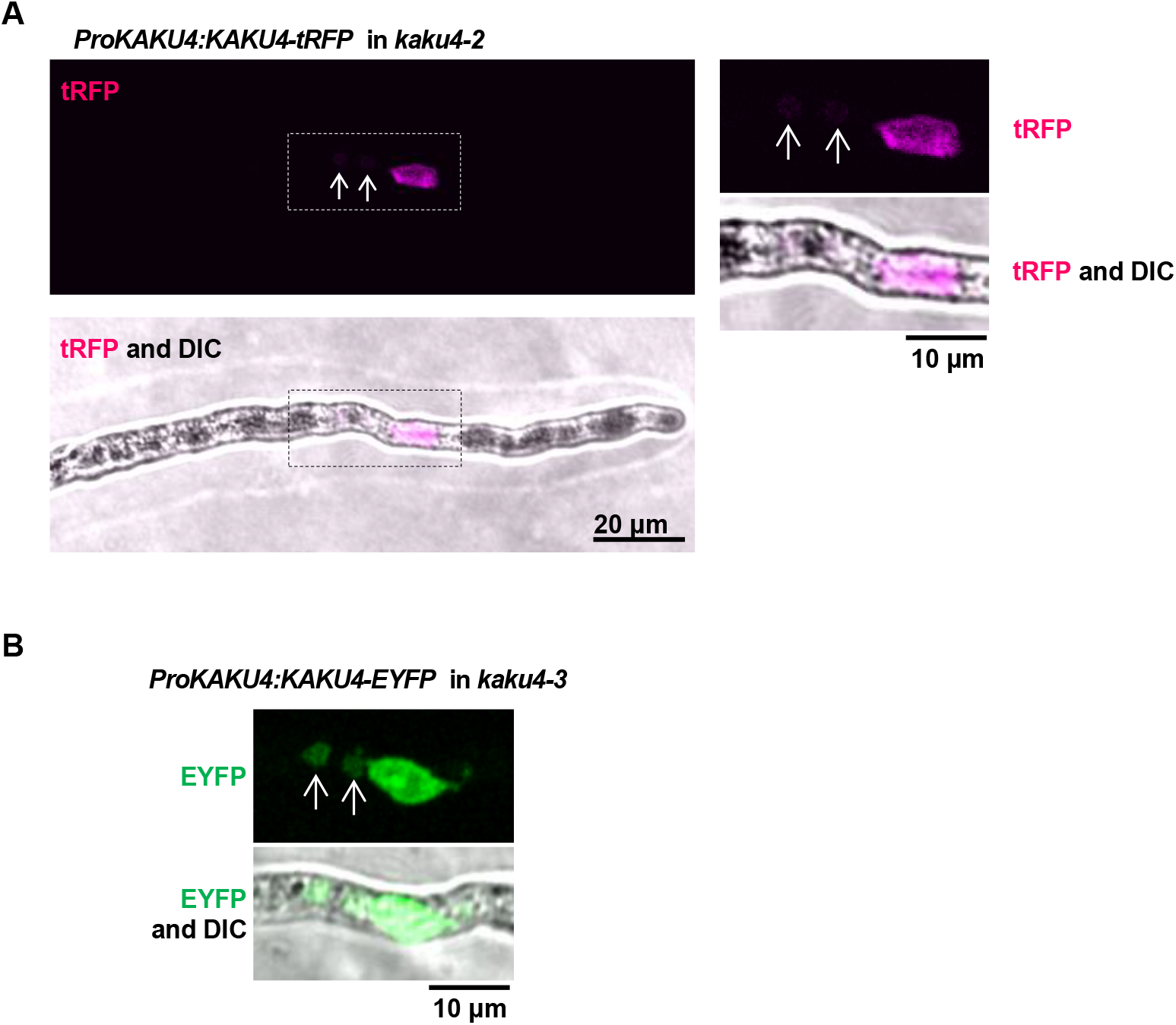
Nuclear envelope localization of KAKU4 in elongating pollen tubes. (A) A pollen tube from *kaku4-2* expressing *KAKU4-tRFP* under the control of *KAKU4* promoter. This image was taken after 3 h of incubation using the *in-vitro* pollen germination system. The areas surrounded by dashed lines are enlarged on the right side. (B) A pollen tube from *kaku4-3* expressing *KAKU4-EYFP* under the control of *KAKU4* promoter. This image was taken after 3.5 h of incubation using the *in-vitro* pollen germination system. The pollen tube elongates toward the right of the panels. Arrows indicate SCNs. DIC, differential interference contrast.

To quantitatively analyze the migration order of the VNs and the SCNs, elongating pollen tubes were categorized into three types: (1) the VN-first type, in which the VN preceded the two SCNs (Fig. 5A, upper), (2) the SCN-first type, in which the two SCNs preceded the VN (Fig. 5A, lower), and (3) the together type, in which VN and SCN were close to each other. Most of the wild type pollen tubes (91–93 %) were VN-first type (Fig. 5B). The proportions of VN-first type were markedly reduced to 33–52% in three *kaku4* mutant alleles (Fig. 5B), indicating that the control of the migration order of the VN and the SCNs in pollen tubes was impaired in the *kaku4* mutant alleles.

**Fig. 5.**
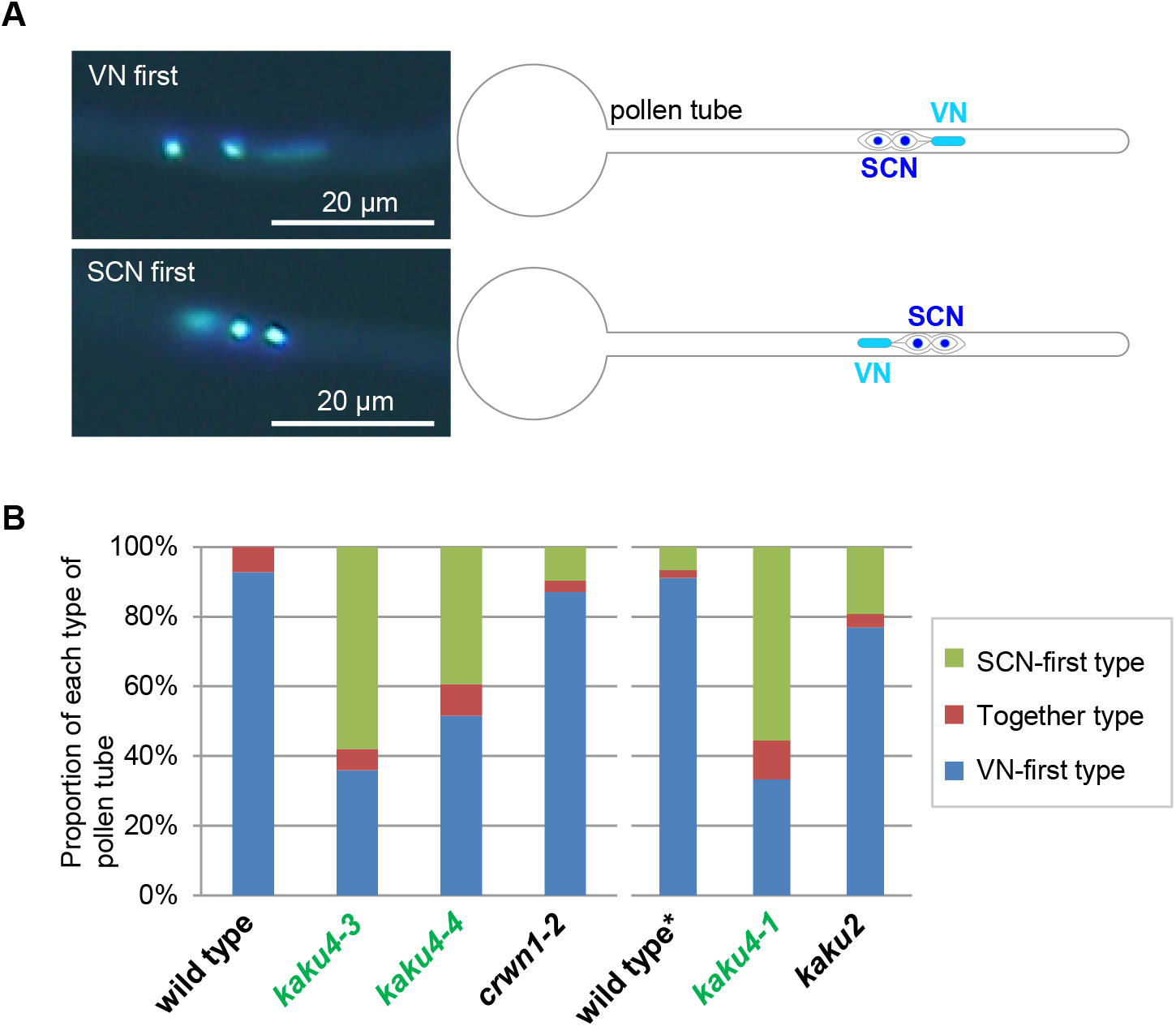
Position of VN and SCN in pollen tubes of *kaku4*. (A) (Left) Representative fluorescence images of nuclei in pollen tubes stained with Hoechst 33342. (Right) Diagrams of the photos showing the VN and two SCNs in pollen tubes. The diagrams are after Sprunck et al. 2014. (B) Relative positions of VN and SCNs in wild-type, *kaku4* or *kaku2/crwn1* pollen tubes. Pollen grains were germinated on artificial medium at 22 °C. The pollen tubes were fixed and stained with Hoechst 33342 after 3 h (for wild type, *kaku4-3*, *kaku4-4*, and *crwn1-2*) or 3.5 h (for wild type*, *kaku4-1* and *kaku2*) of incubation. The relative positions were categorized as VN-first type (VN is ahead of SCN), SCN-first type (SCNs are ahead of VN); and Together type (VN and SCN are close). In total, 55 (wild type), 50 (*kaku4-3*), 66 (*kaku4-4*), 31 (*crwn1-2*), 45 (wild type*), 18 (*kaku4-1*), and 26 (*kaku2 [crwn1]*) pollen tubes were tested. Wild type*, a transgenic plant expressing Nup50a-GFP (parental line of *kaku4-1* and *kaku2*).

We next examined an effect of deficiency of the KAKU4-interacting protein CRWN1 (Dittmer *et al.*, 2007; Sakamoto and Takagi, 2013) on the migration order by using two mutant alleles (*crwn1-2 and kaku2*) (Goto *et al.*, 2014). The proportions of the VN-first type in these mutant alleles (77-87%) were slightly lower than those of the wild type (Fig. 5B). Taken together, these results suggest that the nuclear lamina candidate proteins KAKU4 and CRWN1 control the migration order of the VN and SCNs in elongating pollen tubes, in which KAKU4 functions more predominantly than CRWN1.

We next compared the nuclear shape and size in the elongating pollen tubes between the wild-type and *kaku4-3* plants, by visualizing the VNs and the SCNs with a nuclear marker (H2B-tdTomato) under the constitutive promoter and by visualizing the VNs with another nuclear marker (NLS-Clover) under the VN-specific promoter. The SCNs exhibited moderate difference in the sizes and shapes between the wild type and *kaku4-3* (Fig. 6A and Supplemental Fig. S2). On the contrary, compared with the *kaku4-3* VNs, the wild-type VNs were larger and highly invaginated (Fig. 6A). To quantitatively evaluate the degree of nuclear deformation, we calculated a circularity index (4πA/P^2^; where A = area of nucleus and P = perimeter of nucleus). The VN circularity index was much smaller for the wild-type than it was for *kaku4-3* in both VN- and SCN-first-type pollen tubes (Fig. 6B, upper). This result indicates that the wild-type VNs are more deformed than the *kaku4-3* VNs. In addition, the fluorescence area of the VNs was larger in the wild-type VNs than in *kaku4-3* (Fig. 6B, lower). Notably, the *kaku4-3* VNs of VN-first-type pollen tubes were significantly smaller than those of the SCN-first-type tubes, although the circularity indices of the VNs of both types of pollen tubes were not significantly different (Fig. 6B). This indicates that having a small size makes it easier for the VNs to precede the sperm cells in elongating pollen tubes.

**Fig. 6.**
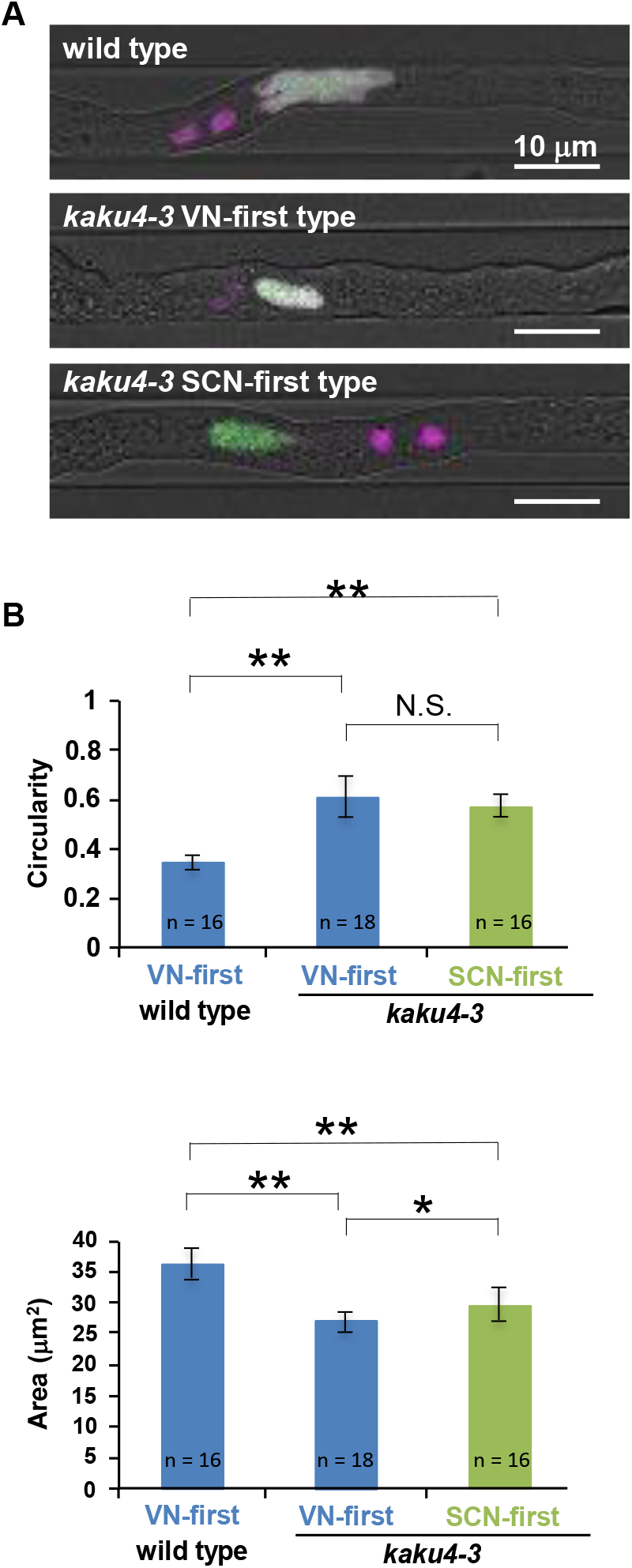
Nuclear shape and size of VN in pollen tubes of *kaku4*. (A) Representative fluorescence images of pollen tubes from the double nuclear marker line, in which the VNs were labeled with H2B-tdTomato (magenta) and NLS-Clover (green) and the SCNs were labeled with H2B-tdTomato (magenta). The pollen tubes were elongated in 10-µm-wide and 4-µm-depth channels of a polydimethylsiloxane microfluidic device and were analyzed after 1 h of incubation at 22 ºC. (B) Circularity indices and areas of the VN in the VN-first-type pollen tubes of the wild-type plants and in the VN- and SCN-first-type pollen tubes of *kaku4-3* plants. Wild-type pollen tubes showed only VN-first-type positioning. The circularity index is defined as the equation 4πA/P^2^ (where A = area of nucleus and P = perimeter of nucleus) and indicates how closely each nucleus corresponds to a spherical shape (a perfect sphere has a circularity index of 1). Any deviation from a circular shape (e.g., elongated, lobulated, or spindle shaped) causes the index to decrease. Means ± standard errors for n = 16 (wild type), 18 (*kaku4-3* VN-first), or 16 (*kaku4-3* SCN-first). Asterisks indicate a significant difference (Student’s *t* test, **P < 0.001, *P < 0.005).

### Slightly decreased seed production and lower pollen competitive ability in *kaku4*

The preceding observations raised the question of whether the migration order of VN and SCN has an effect on fertilization. To assess the effect of *KAKU4* deficiency on fertilization, we analyzed the seed numbers and the silique lengths in the wild type and *kaku4* mutant alleles (Fig. 7A). The number of seeds per silique of the *kaku4* mutant alleles *kaku4-2* and *kaku4-3* was moderately decreased to 82% and 87%, respectively, as compared to that number of the wild type (Fig. 7B). The silique was 1 mm shorter in the *kaku4* mutants, than in wild-type plants (Fig. 7C). These results indicate that *KAKU4* contributed to efficient seed production and proper fruit development.

**Fig. 7.**
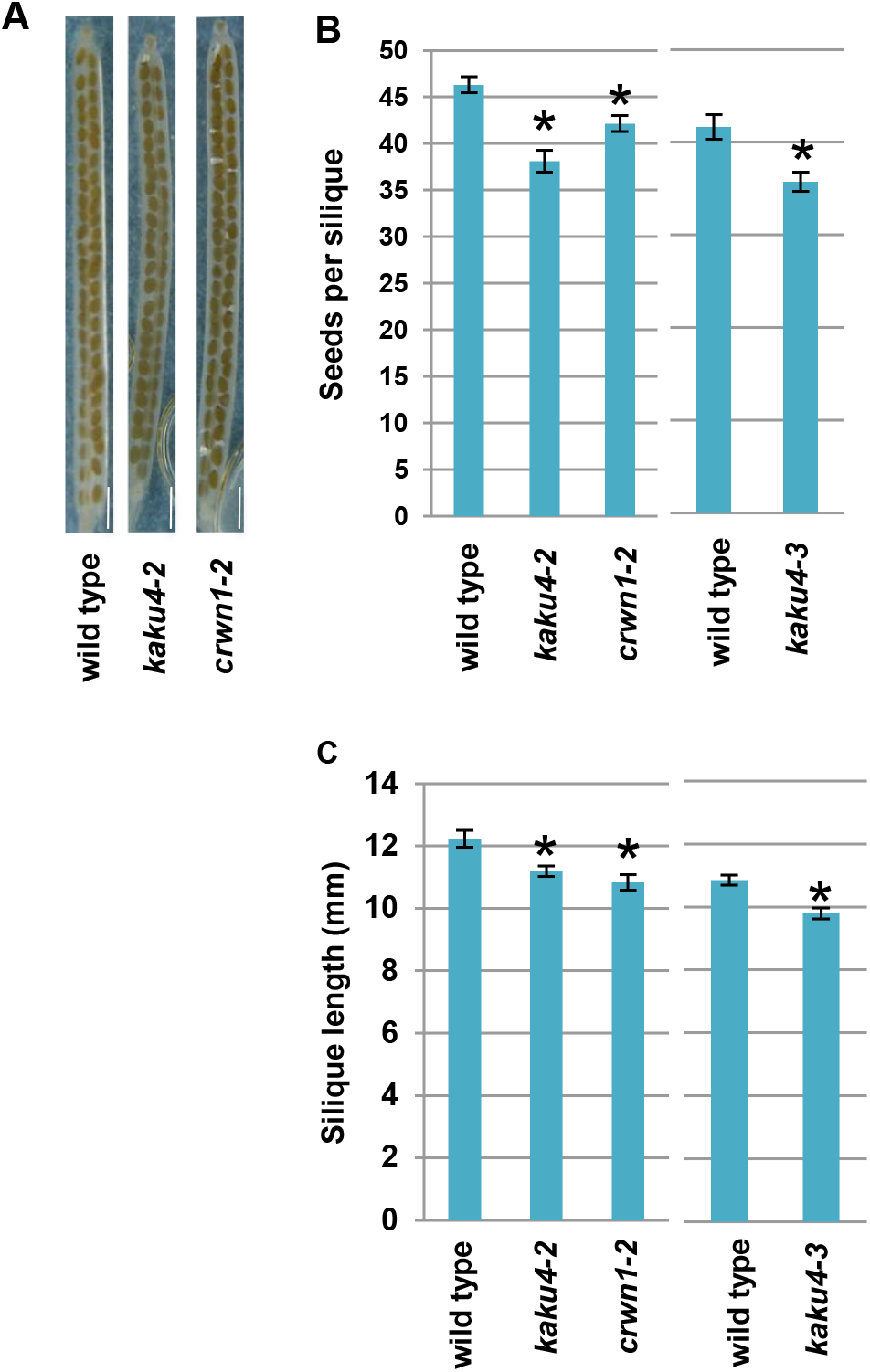
Seed production in *kaku4*. (A) Representative photographs of siliques of wild-type plant, *kaku4-2*, and *crwn1-2*. Bars = 1 mm. (B) Number of seeds per silique from wild-type plant, *kaku4-2*, and *crwn1-2* (means ± standard errors for n = 60) (left) and the number of seeds per silique from wild-type plant and *kaku4-3* (means ± standard errors for n = 59 [wild-type] or 55 [*kaku4-3*]) (right). (C) Length of the siliques from wild-type plant, *kaku4-2*, and *crwn1-2* (means ± standard errors for n = 23 [wild-type], 22 [*kaku4-2*], or 27 [*crwn1-2*]) (left) and length of the siliques from wild-type plant and *kaku4-3* (means ± standard errors for n = 62 [wild-type] or n = 55 [*kaku4-3*]) (right). Asterisks indicate a significant difference from wild type (Student’s *t* test, *P < 0.05).

To test whether the seed reduction in *kaku4* was paternally controlled, we reciprocally crossed between wild-type plants and the heterozygous *kaku4-3*/+ mutant. The *KAKU4* mutant gene in *kaku4-3* carries a kanamycin resistance gene insert. As shown in Table 1, when *kaku4-3*/+ was used as pollen donor to the wild type, the number of kanamycin-resistant progeny was significantly smaller than expected, indicating a reduced pollen competitive ability of *kaku4*. When the wild type was used as the pollen donor to *kaku4-3*/+, the number of kanamycin-resistant progeny did not significantly differ from the expected values. This suggests that *KAKU4* has little or no involvement in functions of pistils in reproduction. Taken together, these results suggest that the VN precedence over SCs in pollen tubes contributes to maintaining proper male transmission efficiency in seed production.

**Table1.** Genetic assay of male transmission in *kaku4-3*/+ mutant by reciprocal crosses.

| Female parent | Male parent | Kan<sup>R</sup> | Kan<sup>S</sup> | Kan<sup>R</sup>/Kan<sup>S</sup> | $\chi^2$ | TE<sub>f</sub> | TE<sub>m</sub> |
| --- | --- | --- | --- | --- | --- | --- | --- |
| *kaku4-3/+* | *kaku4-3/+* | 280 | 84 | 3.33333 | 0.39682 | NA | NA |
| wild type | *kaku4-3/+* | 83 | 145 | 0.57241 | 0.00004 | NA | 57% |
| *kaku4-3/+* | wild type | 126 | 129 | 0.97674 | 0.85098 | 98% | NA |

Kan<sup>R</sup>, kanamycin-resistant; Kan<sup>S</sup>, kanamycin-sensitive; TE, transmission efficiency; TE = Kan<sup>R</sup>/Kan<sup>S</sup> (%); TE<sub>f</sub>, female transmission; TE<sub>m</sub>, male transmission; NA, not applicable.

## Discussion

In wild-type pollen tubes, the migration order of VN and sperm cells is mostly maintained during their migration, although it occasionally changes (Zhou and Meier, 2014). This implies that the initial positions of the VN and sperm cells when they move from the pollen grain to the pollen tube is important for the VN precedent migration over the two-sperm-cell unit that is surrounded by a membrane (Li *et al.*, 2013; Sprunck *et al.*, 2014; Wudick *et al.*, 2018). In wild-type pollen tubes, the VN enters the pollen tube earlier than sperm cells even though it is expected to be larger than the two sperm cells combined (Mcconchie *et al.*, 1987). In this study, *KAKU4* deficiency causes the losses of 1) the irregularity in VN shape and 2) the control of migration order of VN and sperm cells at the same time, suggesting that the VN deformation ability is related to the migration order in the pollen tube. Our observations of wild-type pollen grains showed that VN envelope was deeply invaginated inside the nucleus and that VN shapes were elongated in the pollen tube. These features might allow the VN to easily enter the pollen tube. Hence, we propose that a KAKU4-dependent ability to elongate the VN shapes increases the probability that VN will enter the pollen tube first, resulting in the VN precedent migration over sperm cells in pollen tubes.

Two studies have shown that depolymerization of microtubules can affect the migration order of the VN and sperm cells (Astrom *et al.*, 1995; Heslopharrison *et al.*, 1988). In *Galanthus nivalis* pollen tubes, microtubule depolymerization with colchicine caused a large increase in the “generative cell lead” (referred to as “SCN first” in this study) (Heslopharrison *et al.*, 1988), and in *Nicotiana tabacum* pollen tubes, microtubule depolymerization with oryzalin caused a decrease in pollen tubes harboring VN and an increase in pollen tubes harboring generative cells (sperm cells) (Fig. 5 in Astrom et al., 1995). The loss of two proteins essential for the recruitment of γ-tubulin complexes at microtubule organizing centers (GIP1 and GIP2) alters nuclear shape in root tips (Batzenschlager *et al.*, 2013). These results, together with our data, raises the possibility that microtubule depolymerization may cause a change in nuclear shape, which affects the migration order of VN and sperm cells.

While the VN preceded the sperm cells in the majority of wild-type pollen tubes, the sperm cells preceded the VN in the majority of the pollen tubes in multiple mutant alleles of *wip*, *wit*, *wifi* (Zhou and Meier, 2014), and the *sun* mutant line expressing a dominant-negative gene, in which WIP1 and WIT1 were delocalized from the VN envelope (*Lat52pro::ERS-RFP-SUN2Lm sun1-KO sun2-KD*) (Zhou *et al.*, 2015). These studies indicate that WIPs and WITs are needed to control the migration order of VN and sperm cells in pollen tubes. VNs were reported to be normal in multiple mutants deficient in *WIPs* and/or *WITs* (*wip*, *wit*, and *wifi*) (Zhou and Meier, 2014). However, we cannot exclude the possibility that shape of VN in the mutants’ pollen differs from that in wild-type pollen. The nuclear shape in the leaves and root hairs of the *wip* mutant allele is similar to that in *kaku4* (Zhou *et al.*, 2012). However, because the ratio of “VN first” and “SCN first” is very different between the *wip/wit/wifi* and *kaku4* mutants, the underlying mechanism affecting the migration order of VN and SCNs may be different between WIP/WIT and KAKU4.

Defects in the migration order of VN and sperm cells in *wip/wit/wifi* (Zhou and Meier, 2014) and *kaku4* (this study) caused a decrease in seed reproduction, which suggests that positioning VN first in the pollen tubes is important. In mutants deficient in *WIPs* and/or *WITs*, the pollen tubes frequently failed to properly elongate, to burst at the tube tips, and to result in fertilization (Zhou and Meier, 2014). These observations suggest that the proximity of the VN to the growing pollen tube tip is probably critical for pollen tube reception (Zhou and Meier, 2014). The number of seeds in *kaku4* mutants was ~80% of that in wild-type plants (Fig. 7B). This result suggests that *KAKU4* deficiency does not cause severe defects in pollen tube growth and fertilization. One possible explanation for the impaired competitive ability of pollen in *kaku4* is that the elongation speed of SCN-first-type pollen tubes might be slightly lower than that of VN-first-type pollen tubes. Positioning of VN close to the tip may contribute to efficient elongation of pollen tubes.

## Abbreviations

CRWN: CROWDED NUCLEI
SCN: sperm cell nucleus
SUN: Sad1/UNC84 Homology
VN: vegetative nucleus
WIP: WPP domain–interacting protein
WIT: WPP domain–interacting tail-anchored protein.

## Supplementary data

The following materials are available in the online version of this article.

**Supplementary Fig. S1.** Expression of *KAKU4, WIT1, WIT2, and WIP3*. Transcript levels of *KAKU4*, *WIT1*, *WIT2* and *WIP3* in various tissues. Data were obtained from the resource page of the AtGenExpress project (http://jsp.weigelworld.org/AtGenExpress/resources/) (Schmid et al. 2005). Intensities (absolute values) from select tissues of wild-type plants are shown.

**Supplementary Fig. S2.** Nuclear shape and size of SCN in pollen tubes of *kaku4.* Circularity indices and areas of SCNs in the VN-first-type pollen tubes of the wild-type plants and in the VN- and SCN-first-type pollen tubes of *kaku4-3* plants. Wild-type pollen tubes showed only VN-first-type positioning. The circularity index is defined as the equation 4πA/P^2^ (where A = area of nucleus and P = perimeter of nucleus). Means ± standard errors for n = 16 (wild type), 18 (*kaku4-3* VN-first), or 16 (*kaku4-3* SCN-first). Asterisks indicate a significant difference (Student’s t test, **P < 0.001, *P < 0.005).

## Acknowledgements

We are grateful to Tsuyoshi Nakagawa (Shimane University, Japan) for donating the Gateway vectors; and to the Arabidopsis Biological Resource Center for providing the T-DNA tagged lines of *A. thaliana*. This work was supported by a ‘Specially Promoted Research’ Grant-in-Aid for Scientific Research to I.H-N. (nos. 22000014 and 15H05776), by a Grant-in-Aid for Scientific Research to K.T. (nos. 26711017 and 18K06283), and by a Grant-in-Aid for JSPS Fellows to C.G. (nos. 13J01227 and 17J06391) from the Japan Society for the Promotion of Science (JSPS). This work was also supported by the Human Frontier Science Program to K.T. (RGP0009/2018) from the International Human Frontier Science Program Organization.

## References

Astrom H, Sorri O, Raudaskoski M. 1995. Role of Microtubules in the Movement of the Vegetative Nucleus and Generative Cell in Tobacco Pollen Tubes. Sexual Plant Reproduction 8, 61–69.

Batzenschlager M, Masoud K, Janski N, Houlne G, Herzog E, Evrard JL, Baumberger N, Erhardt M, Nomine Y, Kieffer B, Schmit AC, Chaboute ME. 2013. The GIP gamma-tubulin complex-associated proteins are involved in nuclear architecture in Arabidopsis thaliana. Front Plant Sci 4, 480.

Boavida LC, McCormick S. 2007. Temperature as a determinant factor for increased and reproducible in vitro pollen germination in Arabidopsis thaliana. Plant Journal 52, 570–582.

Borg M, Twell D. 2011. Pollen: Structure and Development. In: eLS. John Wiley & Sons, Ltd: Chichester.

Chen LQ, Lin IW, Qu XQ, Sosso D, McFarlane HE, Londono A, Samuels AL, Frommer WB. 2015. A cascade of sequentially expressed sucrose transporters in the seed coat and endosperm provides nutrition for the Arabidopsis embryo. Plant Cell 27, 607–619.

Clough SJ, Bent AF. 1998. Floral dip: a simplified method for *Agrobacterium*-mediated transformation of *Arabidopsis thaliana*. Plant J 16, 735–743.

Dittmer TA, Stacey NJ, Sugimoto-Shirasu K, Richards EJ. 2007. LITTLE NUCLEI genes affecting nuclear morphology in Arabidopsis thaliana. Plant Cell 19, 2793–2803.

Fiserova J, Kiseleva E, Goldberg MW. 2009. Nuclear envelope and nuclear pore complex structure and organization in tobacco BY-2 cells. Plant J 59, 243–255.

Goto C, Tamura K, Fukao Y, Shimada T, Hara-Nishimura I. 2014. The Novel Nuclear Envelope Protein KAKU4 Modulates Nuclear Morphology in Arabidopsis. Plant Cell 26, 2143–2155.

Hamamura Y, Saito C, Awai C, Kurihara D, Miyawaki A, Nakagawa T, Kanaoka MM, Sasaki N, Nakano A, Berger F, Higashiyama T. 2011. Live-cell imaging reveals the dynamics of two sperm cells during double fertilization in Arabidopsis thaliana. Curr Biol 21, 497–502.

Heslopharrison J, Heslopharrison Y, Cresti M, Tiezzi A, Moscatelli A. 1988. Cytoskeletal Elements, Cell Shaping and Movement in the Angiosperm Pollen-Tube. Journal of Cell Science 91, 49–60.

Kawashima T, Berger F. 2011. Green love talks; cell-cell communication during double fertilization in flowering plants. AoB Plants 2011, plr015.

Li S, Zhou LZ, Feng QN, McCormick S, Zhang Y. 2013. The C-terminal hypervariable domain targets Arabidopsis ROP9 to the invaginated pollen tube plasma membrane. Mol Plant 6, 1362–1364.

Maruyama D, Hamamura Y, Takeuchi H, Susaki D, Nishimaki M, Kurihara D, Kasahara RD, Higashiyama T. 2013. Independent Control by Each Female Gamete Prevents the Attraction of Multiple Pollen Tubes. Developmental Cell 25, 317–323.

Maruyama D, Volz R, Takeuchi H, Mori T, Igawa T, Kurihara D, Kawashima T, Ueda M, Ito M, Umeda M, Nishikawa S, Gross-Hardt R, Higashiyama T. 2015. Rapid Elimination of the Persistent Synergid through a Cell Fusion Mechanism. Cell 161, 907–918.

Mcconchie CA, Russell SD, Dumas C, Tuohy M, Knox RB. 1987. Quantitative Cytology of the Sperm Cells of Brassica-Campestris and Brassica-Oleracea. Planta 170, 446–452.

McCue AD, Cresti M, Feijo JA, Slotkin RK. 2011. Cytoplasmic connection of sperm cells to the pollen vegetative cell nucleus: potential roles of the male germ unit revisited. Journal of Experimental Botany 62, 1621–1631.

Meier I, Richards EJ, Evans DE. 2017. Cell Biology of the Plant Nucleus. Annu Rev Plant Biol 68, 139–172.

Muro K, Matsuura-Tokita K, Tsukamoto R, Kanaoka MM, Ebine K, Higashiyama T, Nakano A, Ueda T. 2018. ANTH domain-containing proteins are required for the pollen tube plasma membrane integrity via recycling ANXUR kinases. Communications Biology 1.

Nakagawa T, Suzuki T, Murata S, Nakamura S, Hino T, Maeo K, Tabata R, Kawai T, Tanaka K, Niwa Y, Watanabe Y, Nakamura K, Kimura T, Ishiguro S. 2007. Improved gateway binary vectors: High-performance vectors for creation of fusion constructs in Transgenic analysis of plants. Bioscience Biotechnology and Biochemistry 71, 2095–2100.

Poulet A, Probst AV, Graumann K, Tatout C, Evans D. 2017. Exploring the evolution of the proteins of the plant nuclear envelope. Nucleus 8, 46–59.

Sakamoto Y, Takagi S. 2013. LITTLE NUCLEI 1 and 4 regulate nuclear morphology in Arabidopsis thaliana. Plant Cell Physiol 54, 622–633.

Schmid M, Davison TS, Henz SR, Pape UJ, Demar M, Vingron M, Scholkopf B, Weigel D, Lohmann JU. 2005. A gene expression map of Arabidopsis thaliana development. Nat Genet 37, 501–506.

Sprunck S, Hackenberg T, Englhart M, Vogler F. 2014. Same same but different: sperm-activating EC1 and ECA1 gametogenesis-related family proteins. Biochem Soc Trans 42, 401–407.

Takeuchi H, Higashiyama T. 2016. Tip-localized receptors control pollen tube growth and LURE sensing in Arabidopsis. Nature 531, 245–248.

Tamura K, Iwabuchi K, Fukao Y, Kondo M, Okamoto K, Ueda H, Nishimura M, Hara-Nishimura I. 2013. Myosin XI-i links the nuclear membrane to the cytoskeleton to control nuclear movement and shape in Arabidopsis. Curr Biol 23, 1776–1781.

Twell D, Yamaguchi J, Mccormick S. 1990. Pollen-Specific Gene-Expression in Transgenic Plants - Coordinate Regulation of 2 Different Tomato Gene Promoters during Microsporogenesis. Development 109, 705–713.

Vogler F, Konrad SSA, Sprunck S. 2015. Knockin’ on pollen’s door: live cell imaging of early polarization events in germinating Arabidopsis pollen. Frontiers in Plant Science 6.

Wang H, Dittmer TA, Richards EJ. 2013. Arabidopsis CROWDED NUCLEI (CRWN) proteins are required for nuclear size control and heterochromatin organization. BMC Plant Biol 13, 200.

Wudick MM, Portes MT, Michard E, Rosas-Santiago P, Lizzio MA, Nunes CO, Campos C, Santa Cruz Damineli D, Carvalho JC, Lima PT, Pantoja O, Feijo JA. 2018. CORNICHON sorting and regulation of GLR channels underlie pollen tube Ca(2+) homeostasis. Science 360, 533–536.

Yanagisawa N, Sugimoto N, Arata H, Higashiyama T, Sato Y. 2017. Capability of tip-growing plant cells to penetrate into extremely narrow gaps. Sci Rep 7, 1403.

Zhou X, Graumann K, Evans DE, Meier I. 2012. Novel plant SUN-KASH bridges are involved in RanGAP anchoring and nuclear shape determination. J Cell Biol 196, 203–211.

Zhou X, Groves NR, Meier I. 2015. SUN anchors pollen WIP-WIT complexes at the vegetative nuclear envelope and is necessary for pollen tube targeting and fertility. Journal of Experimental Botany 66, 7299–7307.

Zhou X, Meier I. 2014. Efficient plant male fertility depends on vegetative nuclear movement mediated by two families of plant outer nuclear membrane proteins. Proceedings of the National Academy of Sciences of the United States of America 111, 11900–11905.

