## Supplemental Figures for "KAKU4-mediated deformation of the vegetative nucleus controls its precedent migration over sperm cells in pollen tubes"

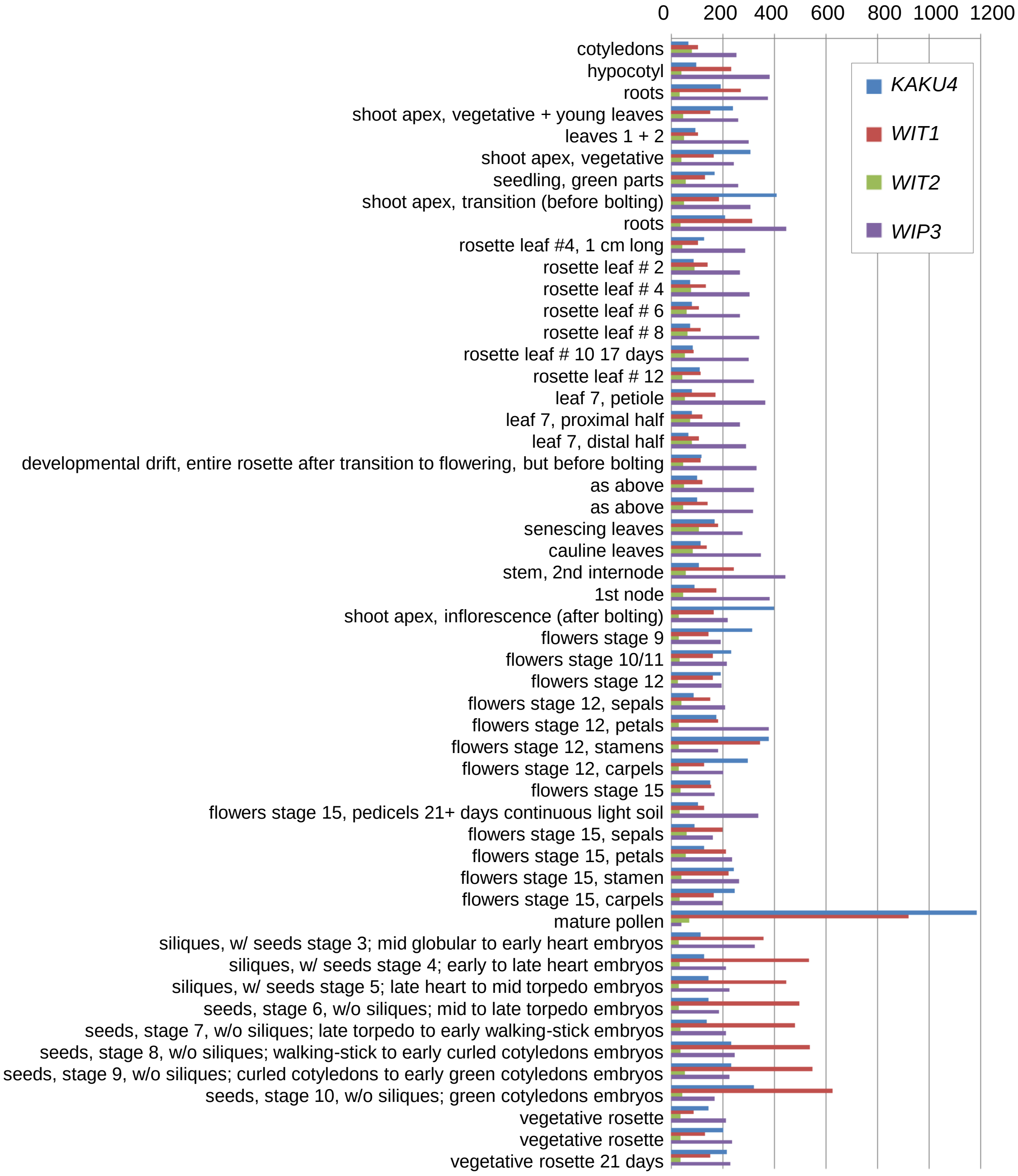


**Supplementary Fig. S1.** Expression of *KAKU4, WIT1, WIT2, and WIP3*. Transcript levels of *KAKU4*, *WIT1*, *WIT2* and *WIP3* in various tissues. Data were obtained from the resource page of the AtGenExpress project (http://jsp.weigelworld.org/AtGenExpress/resources/) (Schmid et al. 2005). Intensities (absolute values) from some tissues of wild-type plants are shown.

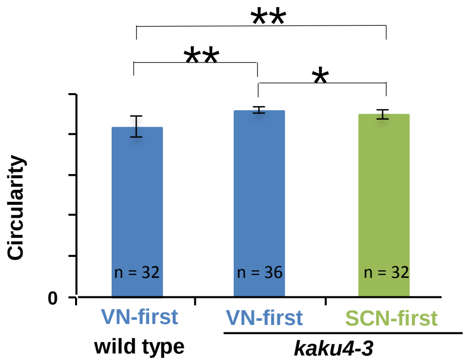


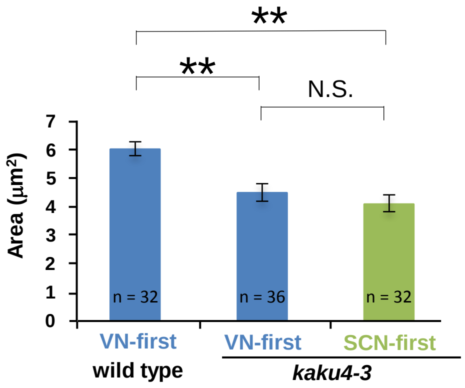


**Supplementary Fig. S2.** Nuclear shape and size of SCN in pollen tubes of *kaku4.*
Circularity indices and areas of SCNs in the VN-first-type pollen tubes of the wild-type plants and in the VN- and SCN-first-type pollen tubes of *kaku4-3* plants. Wild-type pollen tubes showed only VN-first-type positioning. The circularity index is defined as the equation $4\pi A/P^2$ (where A = area of nucleus and P = perimeter of nucleus). Means ± standard errors for n = 16 (wild type), 18 (*kaku4-3* VN-first), or 16 (*kaku4-3* SCN-first). Asterisks indicate a significant difference (Student's t test, $^{**}P < 0.001$, $^{*}P < 0.005$).
